# The kinetics of SARS-CoV-2 nsp7-11 polyprotein processing and impact on complexation with nsp16

**DOI:** 10.1101/2024.01.06.574466

**Authors:** Kira Schamoni-Kast, Boris Krichel, Tomislav Damjanović, Thomas Kierspel, Sibel Toker, Charlotte Uetrecht

## Abstract

In severe-acute respiratory syndrome coronavirus 2 (SARS-CoV-2) infection, polyproteins (pp1a/pp1ab) are processed into non-structural proteins (nsps), which largely form the replication/transcription complex (RTC). The polyprotein processing and complex formation is critical and offers potential therapeutic targets. However, the interplay of polyprotein processing and RTC-assembly are poorly understood. Here, we studied two key aspects: The influence of the pp1a terminal nsp11 on the order of polyprotein processing by viral main protease M^pro^ and the influence of polyprotein processing on core enzyme complex formation. We established a method based on native MS to determine rate constants *k* considering the structural environment. This enabled us to quantify the multi-reaction kinetics of coronavirus polyprotein processing for the first time. Our results serve as a blueprint for other multi-cleavage reactions. Further, it offers a detailed and quantifiable perspective to the dynamic reactions of SARS-CoV-2 polyprotein processing, which is required for development of novel antivirals.

## Introduction

Severe-acute respiratory syndrome coronavirus 2 (SARS-CoV-2) infection can be understood as a series of dynamic molecular mechanisms that lead to multiplication of the viral genome. Two thirds of the ∼30-kb single-stranded (+)-sense RNA genome of SARS-CoV-2 encodes mostly for proteins of the viral replication-transcription complex (RTC) and is directly translated into two overlapping polyproteins (pp1a and pp1ab). The polyprotein internal proteases PL^pro^ (nsp3) and M^pro^ (nsp5) facilitate proteolytic processing of pp1a and pp1ab into mature non-structural proteins (nsps) nsp1-nsp11 and nsp1-nsp16, respectively.^1–3^ The pp1a/pp1ab regions that are cleaved by M^pro^ (nsp5-11 in pp1a and nsp5-16 in pp1ab), are directly involved in viral replication and transcription. While nsp12 and nsp13 harbor the function of the polymerase-helicase complex, the other core enzymes nsp14, nsp15 and nsp16 have indispensable function in RNA modification. nsp14 and nsp16, in complex with nsp10 as their co-factor, play an essential role in the viral life cycle^4^. In higher eukaryotes, the 2’-OH position of the first ribose in an RNA transcript is methylated and helps discriminating between self and non-self RNA transcripts. Thus, activity of nsp10/nsp16 complex as 2’O-methyltransferase facilitates host immune evasion^5^. An overview of the interaction network of nsps can be found elsewhere^6^.

The C-terminal end of pp1a, nsp7 to nsp11, contains four small soluble proteins, which have important regulatory roles in replication, and ends with the thirteen amino acid peptide nsp11 lacking known function. However, the region encoding nsp11 regulates the ratio of pp1a and pp1ab via genomic frameshift^4^. Notably, the efficiency of the frameshift makes the translation of pp1a two to five times more likely than pp1ab, which directly influences the concentration ratio between nsp1-11 (pp1a) and nsp12-16 (only in pp1ab). The excess of regulatory subunits appears relevant for stoichiometric assembly of major protein complexes between nsps from pp1a and pp1ab. One important aspect of subunit concentration is the polyprotein cleavage order, from complete polyprotein into matured nsps, generating heterogenous polyprotein intermediates, here referred to as sub-polyproteins or nsp-precursors. The processing of the polyprotein region nsp7-10 is essential for viral growth^7,8^. Importantly, soluble nsp7-11 sub-polyproteins have been detected in coronavirus infected cells^6^. However, their mechanism of action remains to be elucidated.

The M^pro^ cleavage sequences between SARS-CoV and SARS-CoV-2 are highly conserved^9^. In a previous study, we analyzed the processing of the SARS-CoV polyprotein region nsp7-10 *in vitro*. We observed intermediate processing states, above all, a relatively long-lived nsp7-8 product and a short-lived nsp7-9 product, which informed about the order, with which M^pro^ tackles recognition sites within the folded sub-polyprotein (cleavage sites (cs): cs9/10 > cs8/9 >> cs7/8). The nsp7/8 complex only forms after processing of cs7/8. While M^pro^ cannot cleave the synthetic cs8/9 (P6-P6**′**) peptide, cs8/9 in the sub-polyprotein context is accessible and rapidly processed^10^. This indicated that M^pro^ is additionally regulated by the structural environment of the polyprotein. Further, the formation of the nsp7/8 complex upon processing of nsp7-8-His_6_ sub-polyproteins has been investigated^11^. Notably, distinct assemblies are observed in different coronaviruses despite high sequence and structural conservation^12,13^.

The dynamics of the different processing sites, maturing nsps and the formation of the RTC are not understood in detail. It remains unclear, if the processing order of pp1a is conserved and if it is influenced by nsp11 or cs10/11. Moreover, binding capacity of RTC core enzymes to unmatured polyproteins has not been tested. Here, we aim to investigate the dynamics of *in vitro* processing of SARS-CoV-2 nsp7-11 with native mass spectrometry (MS). Native MS is a highly sensitive method allowing simultaneous monitoring of cleavage products making it very versatile to study dynamic and heterogenous processes^14^. Since in native MS, the natural folding and protein-protein interactions are preserved^15^, the protein complexes formed by the processing products can be detected. By acquiring data at different time points during processing, increasing and decreasing ratios of matured nsps, intermediate processing products and protein complexes are monitored simultaneously. Based on the obtained time-dependent protein ratios, we established a method to extract kinetic rate constants *k* for M^pro^ at the four cs 7/8, 8/9, 9/10 and 10/11. Furthermore, we used different recombinant polyprotein constructs to investigate the influence of the nsp10/11 cleavage site on the processing reaction. In conjunction with modelling and sequence alignment, our results provide a connection between structural environment of the four cleavage sites together with their processing order by M^pro^. Finally, we determine the importance of polyprotein processing on complex formation by performing protein-protein interaction assays between cleaved and uncleaved polyprotein and the methyltransferase nsp16. In summary, our study provides detailed kinetics of nsp7-11 multi-cleavage reaction by M^pro^ and validates a mechanistic model of SARS-CoV-2 polyprotein cleavage and nsp complex formation.

## Results

### Processing products and complexes of fully processed sub-polyprotein

In order to investigate polyprotein processing in SARS-CoV-2 we used the nsp7-11 region, which represents the C-terminal end of pp1a. Therefore, two protein constructs containing a polyhistidine-tag (His_6_) on either the N- or the C-terminus (nsp7-11N and nsp7-11C, respectively) were generated and the influence of His_6_-tag on the structural environment of nsp11 or nsp7 (Figure 1 A) was tested. Upon complete digestion of nsp7-11C and nsp7-11N with M^pro^, we performed native MS and determined cleavage products experimentally in native conditions with sub amino acid mass resolution confirming the four naturally occurring and conserved cleavage sites (cs7/8, cs8/9, cs9/10 and cs10/11). The products from nsp7-11C (60.9 kDa) and nsp7-11N (61.1 kDa) differed only in the presence of the His_6_-tag in either nsp11-His_6_ or His_6_-nsp7, respectively. Furthermore, we observed the heterotetrameric protein complex nsp7/8 (2:2) (62.2 kDa and 65.1 kDa) for either construct (Figure 1 D). Hence, the N-terminal His_6_-tag does not impair complex formation, which confirms proper folding of the proteins. Here upon polyprotein processing, the formation of the heterotetrameric complex of nsp7/8 could be shown despite the heterogenous composition of processing products highlighting the specificity of this complex. The processing products could also be confirmed by SDS-polyacrylamide gel electrophoresis (SDS-PAGE), albeit with lower resolution and without details on non-covalent complexes (Supplementary Figure S1).

**Figure 1.**
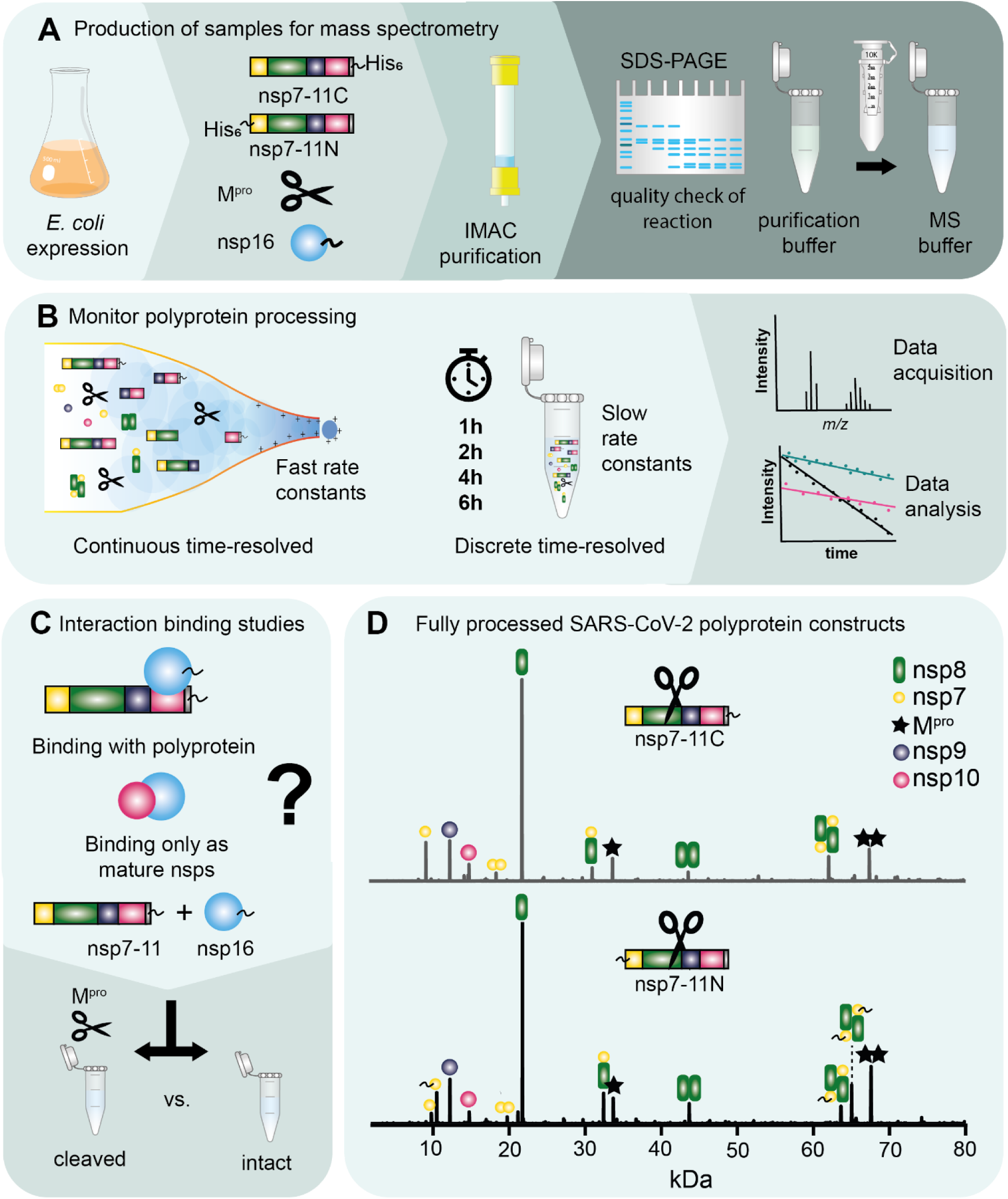
Graphical description of sample preparation and experimental implementation. (A) Four different proteins were recombinantly expressed in *E. coli*. and affinity purified. Prior to MS analysis the samples were exchanged into the volatile MS buffer surrogate ammonium acetate to prevent mass adduct formation. Protein quality was checked by SDS-PAGE. (B) In order to capture fast and slow rate constants of the multi-kinetic cleavage reaction, a two-step measurement approach was required for the analysis, continuous in capillary and discontinuous or discrete time-resolved, respectively. (C) For probing the interaction between the polyprotein and nsp16, M^pro^ was added to process the polyprotein (cleaved) or left untreated (intact). (D) Complete processing of both constructs (nsp7-11C top, nsp7-11N bottom) by M^pro^ (after 20 h incubation) is shown as deconvoluted spectra.

### Continuous monitoring for early cleavage product nsp11

By using native MS, all occurring cleavage products can be detected in parallel and hence be monitored over time. The challenge is to gain sufficient time points when enzymatic reactions happen on different time scales with fast and slow components. Therefore, we conducted continuous time-resolved measurements for faster reactions and discrete time-resolved measurements for slower reactions (Figure 1 B). We started with the continuous time-resolved data acquisition. While time-resolved measurements have been done previously for other samples^16^, having identical starting points is an essential requirement to extract kinetic data of high quality. In order to monitor the fast reactions with early cleavage products, processing was recorded from minute one after start of the reaction, till 30 minutes as last timepoint (Figure 2).

**Figure 2:**
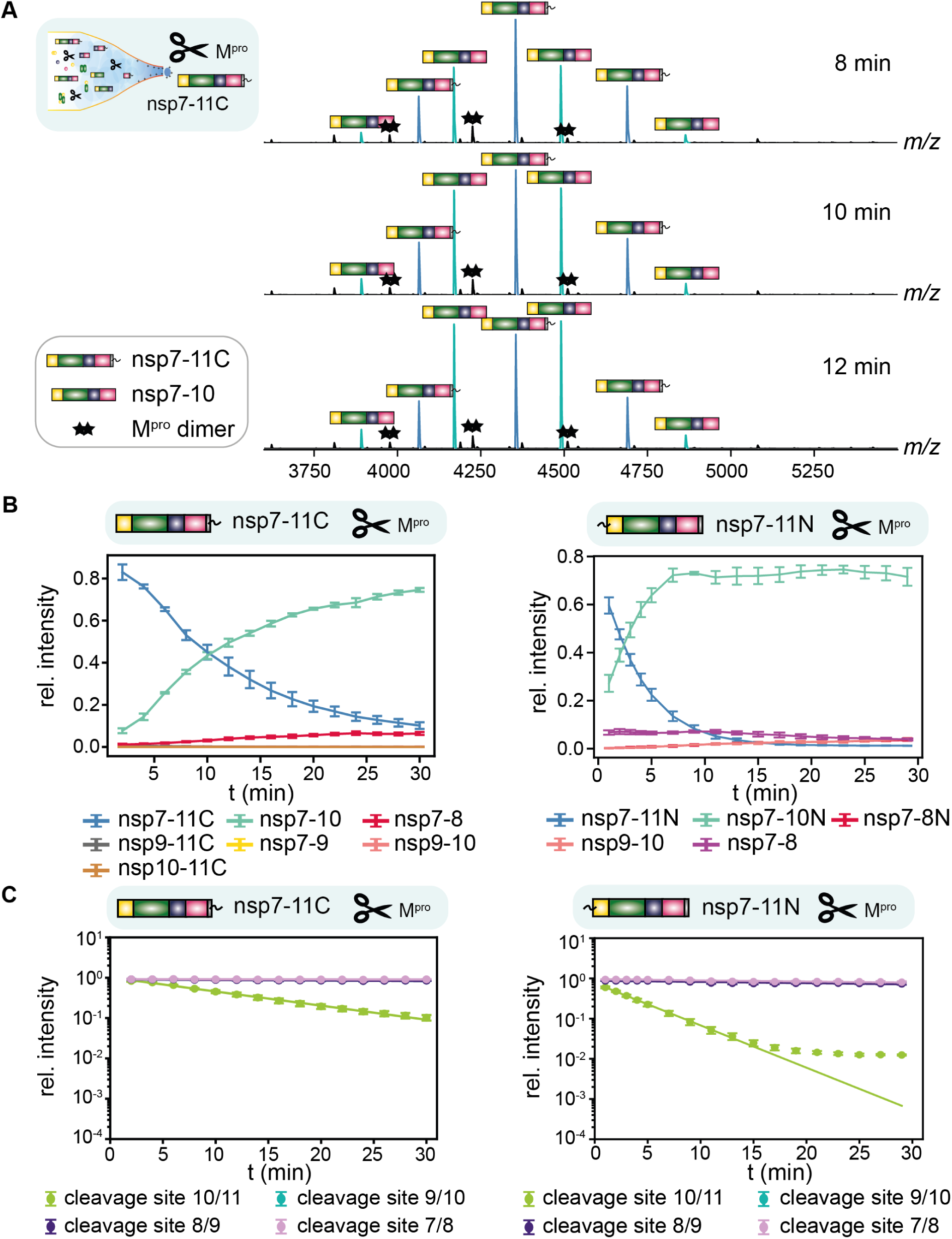
Comparison of continuous processing of nsp7-11C and nsp7-11N. (A) Representative mass spectra of nsp7-11C showing continuous in-capillary processing at 8 min, 10 min and 12 min. (B) Decays of intermediate species of nsp7-11C and nsp7-11N are plotted dominated by nsp7-11C/nsp7-11N (blue) and nsp7-10/nsp7-10N species (turquoise). Data points were simply connected for enhanced visibility. (C) Decline of cleavage sites 10/11 and 9/10 for nsp7-11C and nsp7-11N were plotted on a logarithmic scale and data points were fitted.

With the continuous approach, we observed a decrease of intensity of the polyprotein constructs, nsp7-11C or nsp7-11N, and an increase of the cleavage intermediates nsp7-9 (43.5 kDa), nsp7-10 (58.4 kDa), nsp9-10 (27.3 kDa), nsp9-11 (29.9 kDa) and nsp10-11 (17.5 kDa) in the first 20 min of the processing reaction (Figure 2). This enabled us to monitor the relatively faster cleavage reactions of two processing sites in detail, cs10/11 and cs9/10. cs10/11 in nsp7-11N is released three times faster than in nsp7-11C, indicating that the His_6_-tag decreases cleavage site efficiency. (Figure 2, Table 1). However, nsp11 influences neither the order of processing nor the conversion rates of neighboring cleavage sites. In order to determine cleavage site specific rate constants, the multi-reaction polyprotein processing was simplified to first order kinetics. All detected substrates and intermediates comprising a specific cleavage site were summed up and plotted on a logarithmic scale over time. The data points were then linearly fitted, which showed that simplification was appropriate, and rate constants could be deduced directly from the slope (Figure 2 C). For example, for fitting of the cs10/11 the sub-polyprotein nsp7-11C, nsp9-11 and nsp10-11 were considered (nsp8-11 is not observed). Since the latter two show very low intensities (< 1 %), the rate constant is dominated by the decrease of nsp7-11. Therefore, in the first 30 min, the main conversion is the cleavage of nsp11. The extracted rate constants of cs 9/10, 8/9, 7/8 are similar and relatively slow. In sum, continuous processing indicates that the His_6_-tag affects the speed but not the cleavage order (Table 1, Figure 2 C). Clearly, cleavage of cs10/11 is approximately 48 times faster for nsp7-11N and 14 times faster for nsp7-11C than the others and hence, structurally more accessible for both constructs.

**Table 1.** Cleavage sites with respective mass species are shown with their fitted rate constants in *k* / min^-1^ for the two constructs nsp7-11C and nsp7-11N measured independently, the standard error of the mean (SEM) is also provided.

| Cleavage site | Mass species | $k_{\text{nsp7-11C}} / \text{min}^{-1}$<br>@ 20°C | $k_{\text{nsp7-11N}} / \text{min}^{-1}$<br>@ 20°C |
| --- | --- | --- | --- |
| cs10/11 | nsp7-11C, nsp9-11C<br>nsp10-11C, nsp7-11N | $0.806 \pm 0.002$ | $0.240 \pm 0.005$ |
| cs9/10 | nsp7-11N, nsp7-10N,<br>nsp7-11C, nsp7-10C,<br>nsp9-11C, nsp9-10 | $0.003 \pm 0.004$ | $0.005 \pm 0.001$ |

Ultimately, the continuous processing approach works well for early kinetics, but is time limited due to heating and acidification processes within the capillary. In order to investigate the rate constants *k* at cs8/9 and cs7/8, longer time scales are required. Therefore, we established the discrete time-resolved approach.

### Determining kinetics and ranking the cleavage sites

To monitor slower reactions, we acquired native mass spectra at discrete time-points from 0.5 h to 24 h. Over this long period, a more complete picture emerges. Intermediate cleavage products such as nsp7-10, nsp7-9 nsp7-8 and nsp9-10 increase at early time-points (3-4 h) and then decrease again at later time-points (Figure 3). The intermediate nsp7-10 reached its highest intensity after one hour, while other intermediates only reach their highest intensity after two or more hours before they are gradually converted into mature nsps. As in SARS-CoV^10^, C-terminal cleavage products can be observed early due to more efficient processing at the C-terminal cleavage sites, suggesting a cleavage order from the C- to the N-terminus. However, deviations from this order are observed. For instance, intermediate species such as nsp9-10, nsp10-11 or nsp9-11 were detected, whereby nsp7-9, nsp10-11 and nsp9-11 have low intensities, below 10 %, and are specific to nsp7-11 constructs (Supplementary Figure S2). Notably, there were no intermediate species such as nsp8-9, nsp8-10 or nsp8-11 confirming that cs7/8 cleavage is impaired and occurs last.

**Figure 3:**
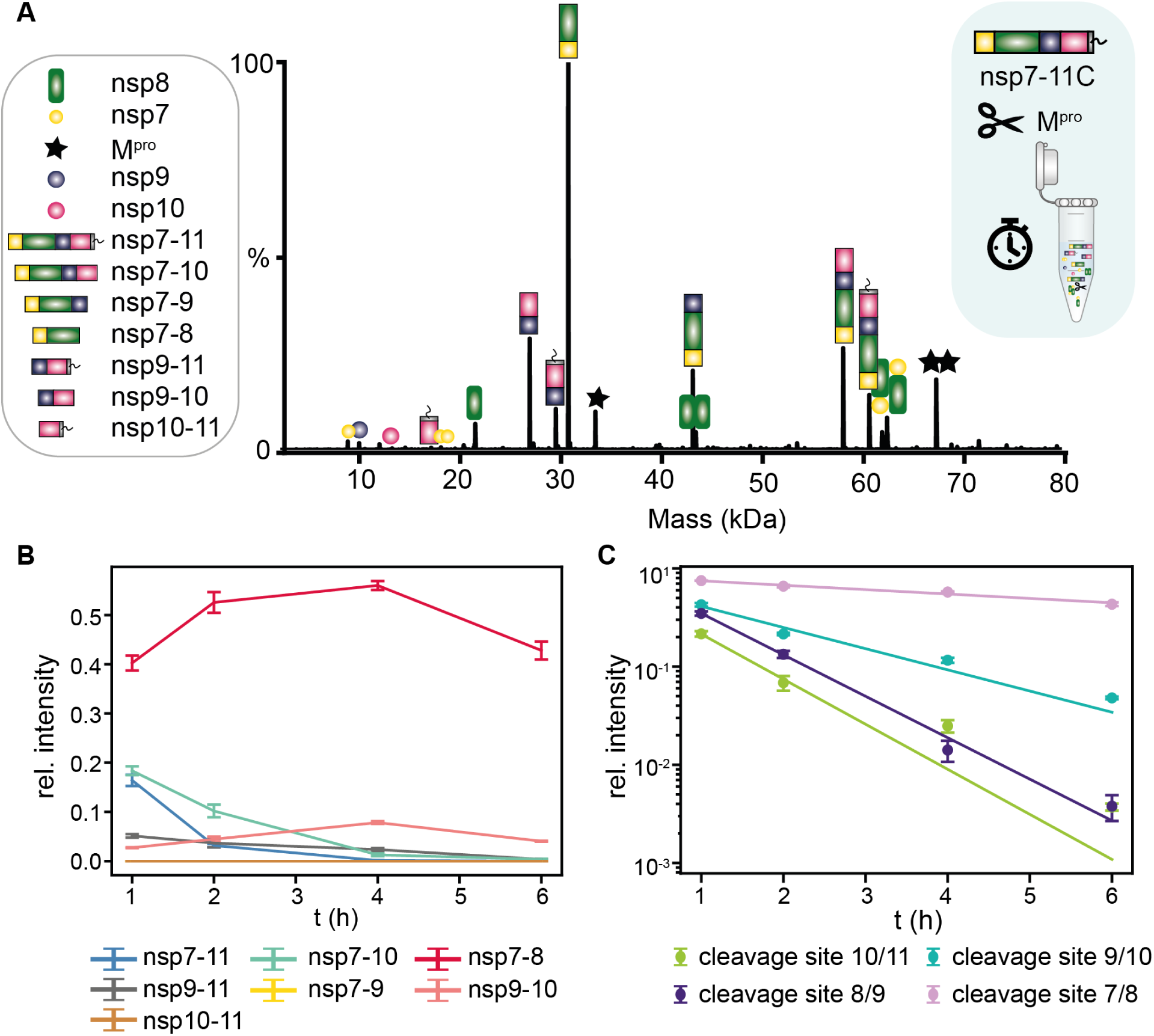
Discrete processing approach of nsp7-11C. (A) A representative deconvoluted mass spectrum after 2h. (B) Course of all deconvoluted intermediate species of the sub-polyprotein at indicated time points. Data points are connected for better visibility. (C) Determination of the rate constants *k* by following the depletion of the substrates corresponding to each cleavage site. The decay is represented as fitted line. After 24 h, all cleavage sites are processed and hence the data points, which are devoid of the plotted species, omitted.

The fitted rate constants *k* at discrete time points reveal the reaction kinetics of the polyprotein processing with cs8/9 and cs10/11 cleaved relatively fast, opposite to the slowest value shown for the cs7/8 (Table 2, Figure 3 C). At *t* > 15 min (nsp7-11N) and supposedly at *t >* 30 min (nsp7-11C) at 20°C in the continuous approach, the data points do not follow an exponential decay anymore due to reduced substrate availability. For nsp7-11C in the discrete approach, this would hence be expected around or after 2 h at 4°C in accordance with our results. Since cs7/8 and cs9/10 have identical amino acid sequence from P2 to P1’^17^, the difference in cleavage kinetics suggests that other mechanisms than the primary sequence are responsible for 7/8 being a less efficient substrate for M^pro^. Intriguingly, we find that cs7/8 is cleaved only when all other sites are processed, suggesting a sterically blockade of the nsp7/8 cleavage site. The second lowest value is fitted for cs9/10 and the occurrence of intermediate species nsp9-11 and nsp9-10 implies a certain preference for cs8/9. Thus, the conducted quantification of the multi-cleavage reaction of pp1a nsp7-11 by M^pro^ results in the cleavage order 10/11 > 8/9 > 9/10 > 7/8.

**Table 2:**
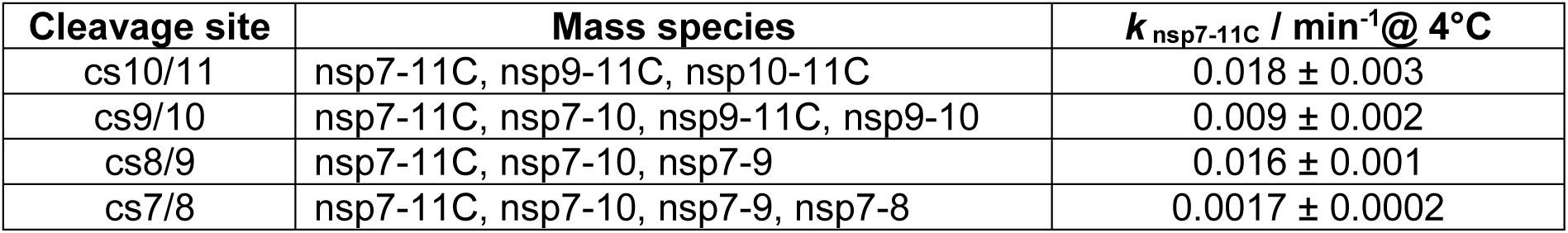
Mass species belonging to the respective cleavage sites are shown with their fitted conversion rates *k* / min^-1^ and the corresponding standard error (SEM). Fitted from processing of nsp7-11C.

| Cleavage site | Mass species | $k_{\text{nsp7-11C}} / \text{min}^{-1} @ 4^{\circ}\text{C}$ |
| --- | --- | --- |
| cs10/11 | nsp7-11C, nsp9-11C, nsp10-11C | $0.018 \pm 0.003$ |
| cs9/10 | nsp7-11C, nsp7-10, nsp9-11C, nsp9-10 | $0.009 \pm 0.002$ |
| cs8/9 | nsp7-11C, nsp7-10, nsp7-9 | $0.016 \pm 0.001$ |
| cs7/8 | nsp7-11C, nsp7-10, nsp7-9, nsp7-8 | $0.0017 \pm 0.0002$ |

### Polyprotein processing as prerequisite for nsp16 complex formation

Polyprotein processing is considered a prerequisite for coronavirus RTC formation, however, the validation of this hypothesis through protein-protein interaction studies has not been performed. The data presented above show that the tertiary structure of the sub-polyprotein influences the processing by M^pro^. Hence, the question arises if it also impacts complexation to RTC enzymes. In previous experiments, we already noted that nsp7/8 complexes require processing^11^. Here, we examined with native MS whether polyprotein processing is required to form the methyltransferase complex nsp16/10 or if nsp16 can bind the immature nsp10 domain within the sub-polyprotein.

First, we mixed nsp7-11C and nsp16 and tested their oligomeric states by native MS (Figure 4). We detected nsp7-11C (61.1 kDa) and nsp16-His_6_ (34.9 kDa) as monomers with similar signal intensity but could not assign signals to a complex between these proteins. It should be noted that the His_6_-tag of nsp16 does not impair the binding to nsp10 as subunit in the polyprotein, since neither C-nor N-terminus are involved in the complexation and tagged nsp16 was shown to form complexes in other structural studies^18^. Then, we initiated processing of polyprotein region nsp7-11C by adding M^pro^ and incubated for 20 h to ensure that the reaction was complete. In the resulting mass spectra, we detected the heterodimeric nsp16/10 (48.4 kDa) protein complex, the monomers of nsp10 (15.0 kDa), nsp16 (33.4 kDa), nsp9 (12.4 kDa) and nsp7 dimers (18.5 kDa), as well as the monomers and dimers of M^pro^ (33.9 kDa and 67.8 kDa, respectively). The putative heterodimeric nsp16/10 complex exhibited a high signal intensity, which indicated a specific binding event. Furthermore, the ion signals assigned to the nsp16/10 complex were validated by collision induced dissociation (CID) (Supplementary Figure S3). Hence, we conclude that immature nsp10 domain as part of the polyprotein does not interact with nsp16. However, if M^pro^ enzymatically processes nsp7-11C and subsequently releases the individual nsps, it leads to the formation of a complex between nsp10 and nsp16 (Figure 4 B). Thus, polyprotein processing is required for methyltransferase nsp16 complex formation.

**Figure 4:**
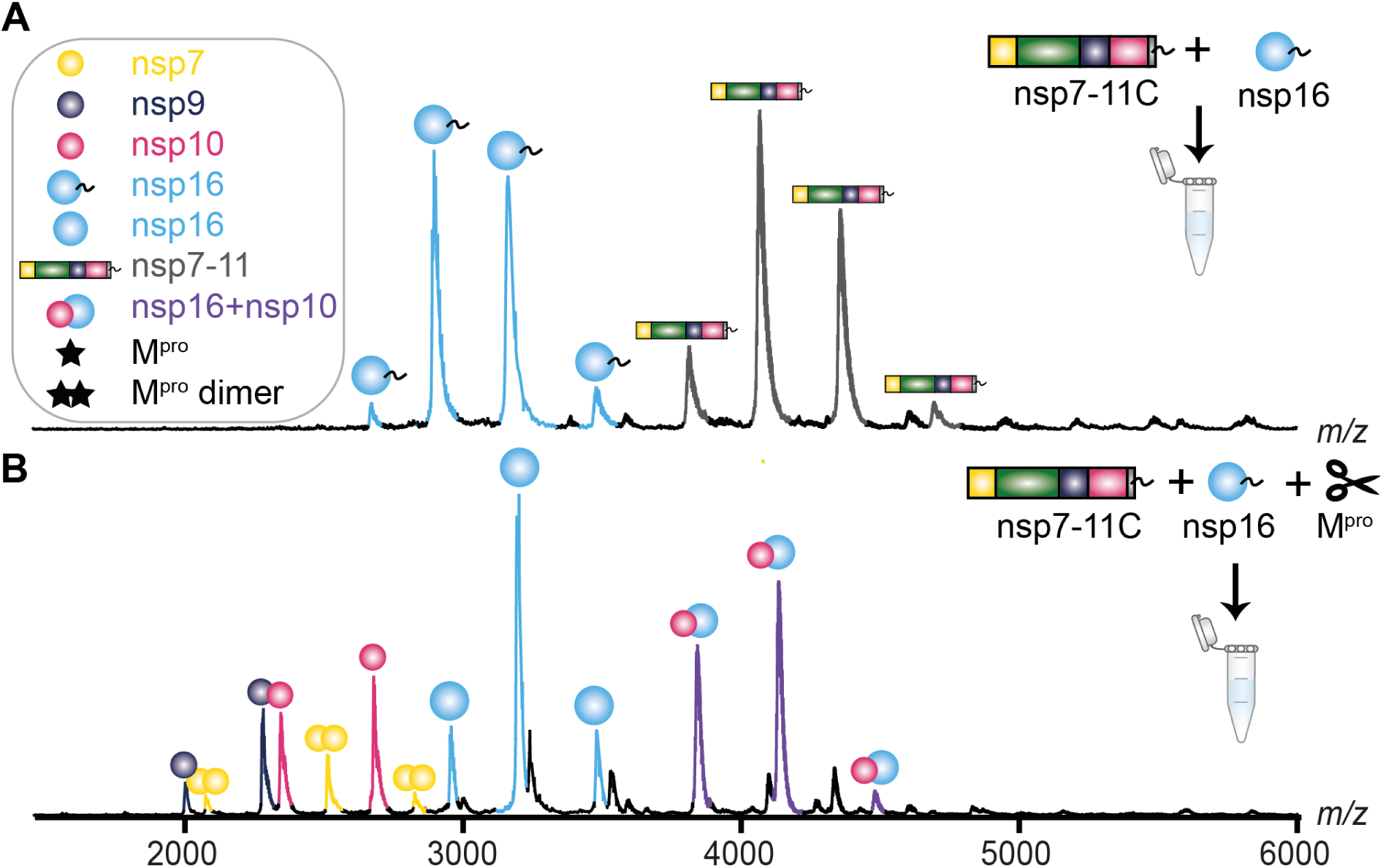
Mass spectra of nsp16 and nsp7-11C. (A) nsp16-His_6_ and nsp7-11C. (B) nsp16-His_6_ is mixed with nsp7-11C and then both are fully processed by M^pro^. Deconvoluted protein species are labeled as indicated on the left panel of the figure. The heterodimeric complex of nsp16/nsp10 appears around *m/z* 4000.

## Discussion

In this work, we have studied the dynamics of polyprotein processing of SARS-CoV-2 by native MS. We show that this method can be used to explore the dynamics of enzymatic processes. For the first time, we have been able to derive rate constants for M^pro^ cleavage sites including the structural environment. By monitoring depletion of all substrate mass species and reducing this process to first-order kinetics, we succeed in quantifying multi-reaction kinetics. We contribute hereby to a better understanding of the regulatory mechanism of polyprotein processing. In the following, we discuss structural implications of our data and later briefly compare our technique to conventional approaches.

The results of our analysis show that in the sub-polyprotein nsp7-11, cs10/11 is the preferred cleavage site for M^pro^, illustrated by the highest determined conversion rate (Tables 1&2). Whereas conversion rates of cleavage sites 8/9 and 9/10 are relatively fast, nsp7-8 accumulates over time, suggesting structural hindrance for M^pro^ to access the 7/8 junction. This is in line with previous observations for SARS-CoV-2 and SARS-CoV^10,21^. Notably, cs9/10 is processed to a lesser extent here than in SARS-CoV. To understand structural hindrance in molecular detail, sequences of the sub-polyprotein and intermediates were submitted to AlphaFold2^22–25^. The predictions resulted in flexible linker regions (plus and minus five amino acids upstream and downstream from the cleavage site, P6-P6’) in nsp8-9, nsp9-10, and nsp10-11. Only the linker region of nsp7-8 was predicted to fold into an α-helix (Figure 5) with high confidence (cf. Supplementary Table S1 and Supplementary Figure S4). A helical shape for this region was also concluded in a previous study^21^. The model and our processing results are in contrast to other predictions showing an unstructured region between nsp7 and nsp8^26^. Therefore, we consider a structured region more likely as helices are poor protease substrates^27^. Furthermore, the flexible linker regions of nsp8-9, and nsp10-11 match the high conversion rates for these cleavage sites. However, the underlying mechanism by which the cleavage order of these faster cleavage sites is determined remains poorly understood.

**Figure 5:**
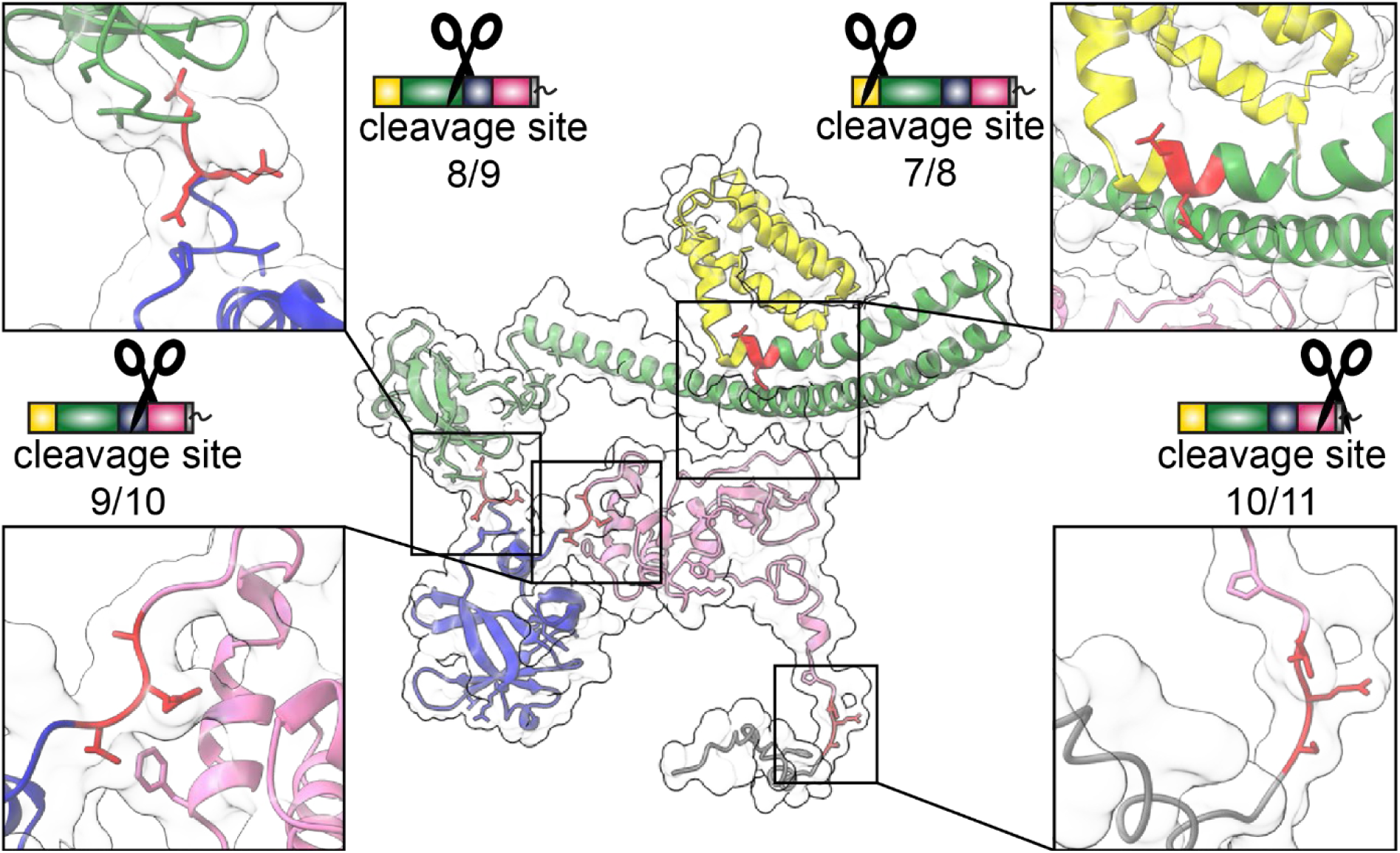
Structure prediction by Alphafold2^38^ of nsp7-11C sub-polyprotein. nsp7 (yellow), nsp8 (green), nsp10 (hotpink) nsp9 (blue) and nsp11-His_6_ (gray). Zoom on cleavage sites (red).

Cleavage efficiency is also influenced by sequence and shows the highest efficiency for LQ/S, which is the interdomain junction from the autocleavage site of M^pro^ and fits to the data showing 10/11, which shares this sequence (LQ/S, P2-P1’), as the fastest cleavage site^19,28,29^. Notably, 7/8 and 9/10 junctions also share the same sequence upfront of the cleavage site (LQ/A, P2-P1’), while showing significantly different rate constants despite the fact that P1’ is less critical. For cs7/8, secondary structure seems to be the key determinant. The intermediate processing speed of cleavage site 9/10 is however not explained by the sequence or the predicted flexible structure. It could be a result of suboptimal residues (P6-P6’) lining the cleavage site, which have been shown to influence cleavage^10^, although the overall influence of the primary structure in folded proteins is low^30^. Narwal et al. proposed two models based on their findings and other studies^11,21^, the polyprotein- and the M^pro^-driven model, in which either the polyprotein or M^pro^ is the key actor^30^. The delayed conversion of cs7/8 is striking, whereas cs9/10 is cleaved at intermediate rates, and cs8/9 as well as cs10/11 are cleaved fast. The overall picture shows a designated order of processing at which the intermediate sub-polyprotein species seem to develop their individual dynamics. They appear to undergo minor conformational changes upon cleavage^21^. Therefore, our data support a polyprotein-driven processing model^30^, which reinforces the question whether the order of processing is conserved among different species and genera of coronaviruses.

In addition to the order of polyprotein processing and its quantification, we also intended to investigate the potential influence of the pp1a terminal nsp11 on the order of processing. The rate constants determined here effectively show that cleavage site 10/11 is the initial section. A C-terminal His_6_-tag further improves accessibility and hence accelerates processing but does not alter processing order. Since nsp11 is cleaved off mostly immediately resulting in the new substrate nsp7-10, we can exclude a major impact of nsp11 on processing of the other cleavage sites. The initial cs10/11 cleavage is contrary to Yadav et al.^21^, who suggested 9/10 as an initial cleavage site using SDS-PAGE and pulsed labeling HDX-MS. While the latter approach offers an elegant observation of distinct dynamic protein populations that are present in solution at certain time points, both methods lack high resolution to distinguish between species. A strict order of processing for the nsp7-11 region is unlikely, as evidenced by species such as nsp9-10, nsp9-11 and nsp10-11. Whereas nsp9-11 and nsp10-11 occurred at rather low intensities (below 5 %), nsp9-10 is more abundant (up to 10 %) and detected for prolonged times. Considering the fitted values for the conversion rates, following cleavage order can be concluded: 10/11 > 8/9 > 9/10 > 7/8. Notably, deleting nsp domains or cleavage sites in the polyprotein was mostly lethal in MHV (Mouse Hepatitis Virus), a β-coronavirus like SARS-CoV-2 with the exception of disruption of cleavage site 9/10 resulting in an attenuated virus^7,8^. This underlines a certain flexibility for processing cleavage site 9/10. However in SARS-CoV, intermediates such as nsp9-10 were not detected and the processing order is from C- to N-terminus (9/10 > 8/9 > 7/8). In contrast to MERS-CoV, in which also alternative intermediates such as nsp9-10 or nsp10-11 seem to occur.^10,11^ Mutations in the polyprotein sequence would be a likely explanation, however sequence and structural alignments did not reveal positions explaining the observed processing differences. Since the substrate specificities (P2-P1’) of M^pro^ among different coronavirus genera are conserved^9^, we conclude that observed differences in processing order point towards structural differences in the polyproteins and potentially distinct regulation.

Conventional techniques often use short labelled peptides as substrates in biochemical assays to demonstrate cleavage efficiency. In this work, we showed an approach how to study and quantify cleavage efficiency including the structure. The here determined *k*-values are much lower even if temperature is considered than *k_cat_* usually measured from biochemical assays in SARS-CoV. Neglecting that SARS-CoV is another strain, the main reason for this is that the peptide substrates are usually added in excess, and therefore the enzyme is always fully occupied. M^pro^ is part of pp1ab, released by autocleavage and then becomes active as a dimer resulting in a twofold excess of the substrate *in vivo*^34^. Thus, the competition of different cleavage sites for the protease active site and similar concentrations of substrate and enzyme reflect the *in vivo* situation much better. Rate constants determined by FRET-assays for SARS-CoV and MERS-CoV show high conversion rates for 7/8 and would thereby propose a different order of processing.^6,20^ Here we showed the lowest rate constant for 7/8. Further in Krichel et al.^10^, the cleavage site 8/9 as a peptide could not be cleaved, but was efficiently processed in the nsp7-10 polyprotein. Cleavage site 8/9 is inherent in the N-terminal NNE sequence of nsp9, which is a suboptimal canonical cleavage site for M^pro^ but important for nsp9 NMPylation function^35^. M^pro^ recognizes non-canonical cleavage sites of Q_(S/A/G/N) (position P1-P1’)^6,36^. Therefore, it is important to include the structural context and probe the cleavage in a more native environment.

While polyprotein processing is key for the formation of the RTC, it remains unclear if certain sub-polyproteins or exclusively mature nsps integrate. Among others, nsp10 and nsp16 need to associate in a functional RTC^8,5,39^ Here, we clearly show that processing is a prerequisite for nsp16/10 complex formation. This result suggests a pathway where a timely release of nsps is essential and highlights the importance of polyprotein processing dynamics.. Such coordinated processing in space and time for regulatory purposes creates a multitude of proteoforms with potential distinct function and is a common pattern in viruses^41,42^. The order and presumed speed regulation of polyprotein processing show parallels to cleavage processes in alphaviruses, which like coronaviruses are enveloped positive sense RNA viruses^40^. In alphaviruses, timeliness and correct processing of ns-polyprotein are crucial for viral replication but are not exclusively dependent on accessibility and sequence.^43,46^

The question arises whether such an intricate coordination of processing also applies to coronaviruses. Genetic experiments have also shown the importance of polyprotein processing (pp1a) for coronaviral replication.^7,8^ The RNA dependent RNA polymerase (RdRp), nsp12, and all RTC enzymes reside in pp1ab and thus occurs in lower concentrations than its binding partners from pp1a such as nsp7, nsp8, nsp9 and nsp10. Only with its processivity factors, nsp7 and nsp8, nsp12 attains its full elongation speed^44^. Further, nsp9 and the kinase-like nidovirus RdRp-associated nucleotidyltransferase (NiRAN) domain from nsp12^45^ execute the essential mRNA capping. The early liberation of nsp9 together with the delayed release of processivity factors could indicate an assembly pathway of the RTC, in which the capping function is established prior to speedy RNA prolongation. Similarly, early release of nsp10 further ensures complexation with nsp14 and nsp16 before nsp12 speeds up, such that they can carry out their functions alongside RNA production. Furthermore, a predicted interaction of nsp9 and nsp8 could not be shown after processing of the nsp7-11 region, which may indicate that nsp12, as the other central interactor, is required as a mediator.^5^ Thus, our study highlights the dependence of processing order on secondary and tertiary structure and shows clear preferences for cleavage sites, which are conceivable as one layer of regulation in RTC assembly.

## Conclusions

In summary, we analyze the proteolytic processing of the C-terminal end of SARS-CoV-2 pp1a, nsp7-11, *in vitro* by the cognate M^pro^ and use native MS to monitor cleavage and complex formation in a time-resolved manner. For the first time, we determined kinetic rate constants *k* for an enzymatic processing by native MS making it an appealing and versatile tool to investigate enzyme kinetics. When combined with commercially available automated injection systems, it lends itself as a complementary screening technique. Assets over conventional techniques are consideration of structural context and label-free substrates that are cheap to produce. A further advantage is the direct feedback on complex formation and stoichiometric ratios in one experiment. Employing temperature control, Arrhenius plots and hence enthalpic and entropic information can be obtained, too^37^.

Furthermore, no structural influence of nsp11 on the processing order is observed. nsp9 and nsp10 are released way earlier than nsp7 and nsp8. Employing Alphfold2, the retarded cleavage of nsp7 and nsp8 likely originates from the α-helical linker. Moreover, nsp16 exclusively binds to processed nsp7-11 providing evidence that processing is a prerequisite for RTC formation. Processing order and complexation with mature nsps show mechanistic potential as regulatory layers coordinating RTC assembly and distinct functions.

## Supporting information

Complete supplement

## Material and Methods

### Production of recombinant proteins

Gene sequences for nsp7-11 and M^pro^ used were taken from “Severe acute respiratory syndrome coronavirus 2 isolate Wuhan-Hu-1” as published in January 2020 (replaced by NCBI LOCUS NC_045512) and commercially synthesized (GenScript). The synthetic gene sequence for nsp7-11C and nsp7-11N with suitable overhangs were cloned with Type IIS restriction enzymes into either pASK35+ and pASK33+ (IBA life sciences), generating a plasmid with C- and N-terminal His_6_-tag, respectively. For expression and purification of nsp7-11 constructs, transformed BL21 Rosetta2 (Merck Millipore) were grown to OD_600_ 0.4-0.6 in 2xYT medium and then induced with 50 µM anhydrotetracycline for 16 h at 20°C. nsp7-11N and nsp7-11C proteins were purified via Ni-NTA affinity chromatography and Superdex10/300 (Cytiva) size exclusion chromatography.^10^ The plasmid for nsp16 was synthesized as full-length nsp16 with N-terminal His_6_-tag in pET22b (+) vector (Supplementary Table S1). For nsp16 expression, the plasmid was transformed in BL21 Rosetta2. Cells were grown until an OD_600_ of 0.4-0.6, cooled on ice and induced with 0.5 M IPTG and then incubated overnight at 20°C. nsp16 was purified via Ni-NTA affinity chromatography and Superdex10/300 (Cytiva) size exclusion chromatography. The plasmid for SARS-CoV-2 M^pro^ in PGEX-6p-1 was generously provided by Prof. Rolf Hilgenfeld, and expressed and purified as described earlier^34,47^.

### Quality control of processing with SDS-PAGE

SDS-Page was performed with a 4%-12% gradient acrylamide Bis-tris gel with XT MES running buffer (Bio-Rad Laboratories). Both constructs nsp7-11C and nsp7-11N were mixed at 36 µM with 14 µM M^pro^ and incubated at 4°C. Aliquots were withdrawn at indicated time points and mixed with XT sample buffers to quench the reaction.

### Native mass spectrometry

Purified samples were exchanged into a structure preserving and MS compatible buffer surrogate ammonium acetate at 300 mM AmAc (99.99 % purity, Sigma-Aldrich), 1 mM DTT, pH 8.0. M^pro^ was exchanged into the buffer surrogate by applying two cycles of centrifugal gel filtration (Biospin mini columns, 6000 MWCO, Bio-Rad). nsp16 and nsp7-11 were exchanged by six rounds of dilution and concentration in centrifugal filter units (Amicon Ultra, 10 K MWCO, Merck Millipore).

A micropipette puller (P-1000, Sutter Instruments) was used to produce nanoESI capillaries in a two-step program from borosilicate capillaries (1.2 mm and 0.68 mm outer and inner diameter, respectively, *World Precision Instruments*) using a squared-box filament (2.5 mm x 2.5 mm). Capillaries were gold-coated by using a sputter coater (CCU-010, Safematic, 5.0 x 10^-2^ mbar, 30.0 mA, 120 s, three runs to vacuum limit 3.0 x 10^-2^ mbar argon). Capillaries were opened using tweezers under a microscope.

For the processing experiments, native MS was performed on a Q Exactive UHMR Orbitrap from Thermo Scientific. Positive ion mode was used by applying capillary voltages of 1.2-1.4 kV, 150°C capillary temperature, 15 V in-source CID and 25 V in HCD cell. Trapping gas pressure optimization was set to 7. Detector optimization was set to “*low m/z*” and the ion transfer *m/z* optimization were adapted as following: Inj. Fl. RF Ampl. to 300, Bent. Fl. RF Ampl. to 940, Trans. MP and HCD-cell RF Ampl. to 900 and C-Trap Ampl. to 2750. Tandem MS were always conducted by the stepwise increase of HCD voltage of 10-20 V.

For the continuous approach, M^pro^ was added to a final concentration of 3 µM to nsp7-11 (final concentration 18 µM), then the sample was briefly mixed by pipetting before transferring 1-2 µL to the capillary. Data acquisition was started 1 min after mixing. At least three replicates were conducted.

For the discrete approach, nsp7-11 and M^pro^ were mixed with a final concentration of 20 µM and 10 µM, respectively. The mixture was then incubated on ice and triplicate measurements were taken at selected time points.

The binding studies of nsp16 and nsp7-11C were implemented on a Q-TOF II mass spectrometer (Micromass/Waters) modified for high-mass experiments (MS Vision)^48^. Final sample concentrations for the interaction studies were 10.2 µM of nsp16 and 12.4 µM of nsp7-11C. For improved electrospray conditions and peak resolution, the sample was diluted to 2.4 µM and 0.4 µM (Figure 4 B) immediately before data acquisition. MS measurements were conducted by using a capillary voltage of 1.3 kV, cone voltage of 130 V and collision cell voltage of 10 V. Pressure in the source region was 10 mbar and argon in the collision cell was adjusted to 1.3 x 10^-2^ mbar. During ESI-MS overview spectra, the MS quadrupole profile was set to 1,000-10,000 *m/z*. The complex of nsp16 and nsp10 was identified by tandem MS by increasing the collision voltage up to 180 V.

### Data analysis

Native mass spectra were investigated and deconvolution was supported by Unidec^49^ (Supplementary Table S3 and Supplementary Table S4). Deconvoluted peaks were checked and *m/z* ranges (gates) were noted to feed into a home-made python script (Supplementary Table S5, Supplementary Table S6). Every gate was plotted and checked before the area under the curve of the detected mass species was taken, which is here called intensity. The initial substrate includes five domains, resulting in five mature proteins. Thus, species intensities were normalized by using a multiplication factor corresponding to the domains or units depending on intermediate species or mature nsps. Since nsp11 was not detected, it was not considered in multiplications.

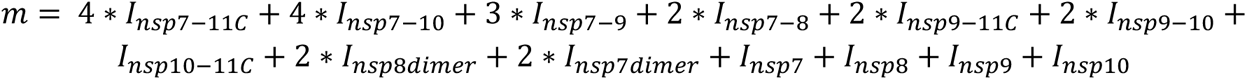

Compared to nsp7-11C, nsp7-11N experiments showed a slightly different composition of detected species. The multiplication array was adapted accordingly:

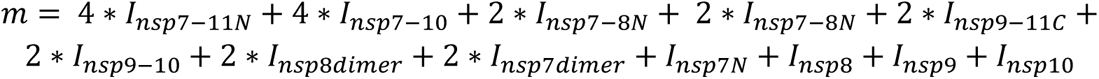

To correct for variation of the spray, the ratio of each individual species to the sum of all species was taken. For example, ratio nsp7-11C including the multiplication:

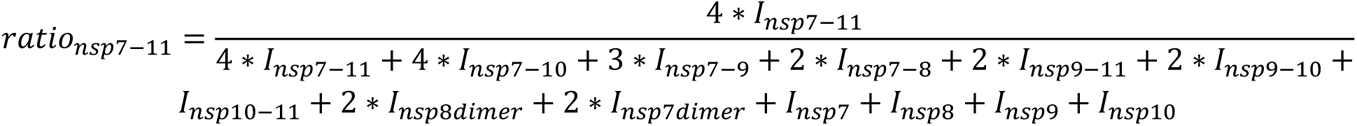

Then these normalized intensities of the replicates were averaged and the standard error of the mean calculated. The fitted rates for the cleavage sites were calculated by using the normalized intensities and summing the species to the corresponding cleavage sites as follows:

Cleavage site 10/11

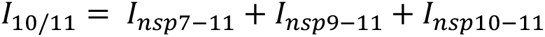

Cleavage site 9/10

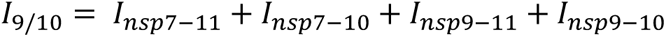

Cleavage site 8/9

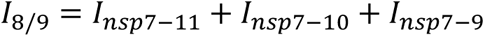

Cleavage site 7/8

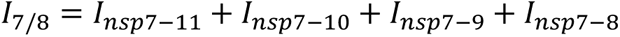

Calculated time-dependent intensities for a given cleavage site were fitted to an integrated form of the first-order kinetics formula:

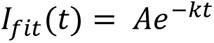

For nsp7-11N, the species were adjusted according to their occurrence. Further, rate constants *k* for each cleavage were extracted from the fit.

### Alphafold2 Modeling

Polyprotein sequences of nsp7-11 (with and without His_6_-tag) and sequences of all intermediates were submitted to Alphafold2 standard run (20 cycles). All models were examined and best models were picked for comparison. Here, the best model for nsp7-11C was selected. pLDDT confidence scores were displayed by using the B-factor column of the pdb-output file. Set color key thresholds in ChimeraX were 50 to 90. Regions with pLDDT measures higher than 70 are expected to be well predicted. Areas that show values between 50 and 70 have low confidence and should be interpreted with caution^38^.

