## Supplementary material for "The kinetics of SARS-CoV-2 nsp7-11 polyprotein processing and impact on complexation with nsp16": Complete supplement

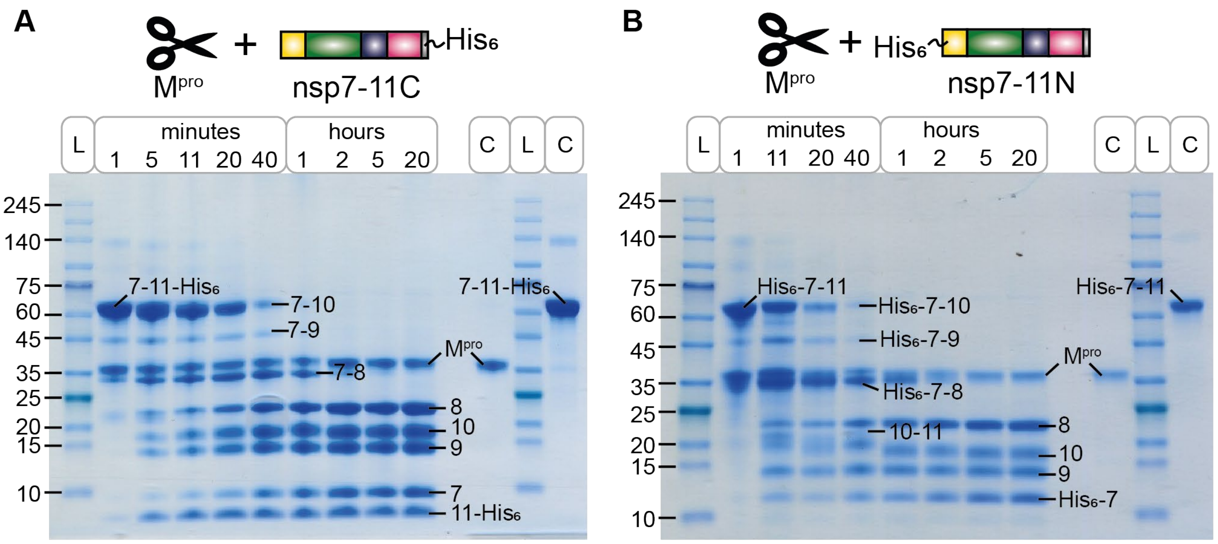


**Supplementary Figure S1: SDS-PAGE of nsp7-11C processing (A) and nsp7-11N processing (B)** showing protein marker ladder (L) and controls (C) of M$^{pro}$ and nsp7-11C or nsp7-11N.

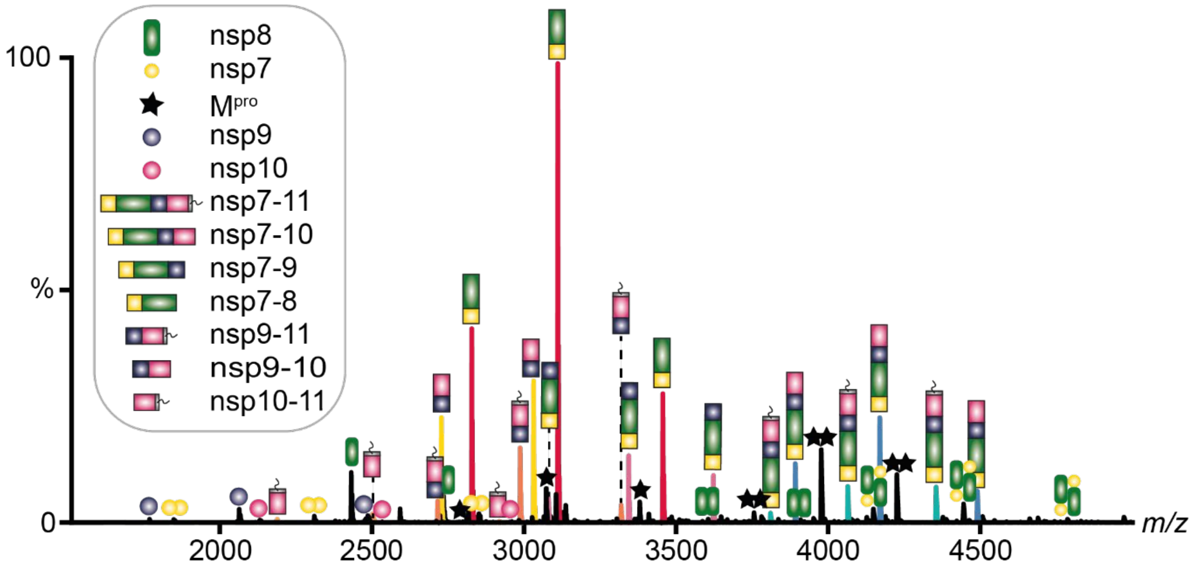


**Supplementary Figure S2: Discontinuous time resolved native MS of nsp7-11C after 2 h.** This is the corresponding mass spectrum to the deconvoluted spectrum (Figure 3 A). Exemplary spectrum of intermediate species. Mature nsps such as nsp8 (21.8 kDa), nsp7 (9.2 kDa), nsp9 (12.4 kDa) and nsp10 (14.9 kDa) are labeled corresponding to their stoichiometry. $M^{pro}$ occurs as monomer (33.8 kDa) and dimer (67.6 kDa) and is marked with a star or double star, respectively.

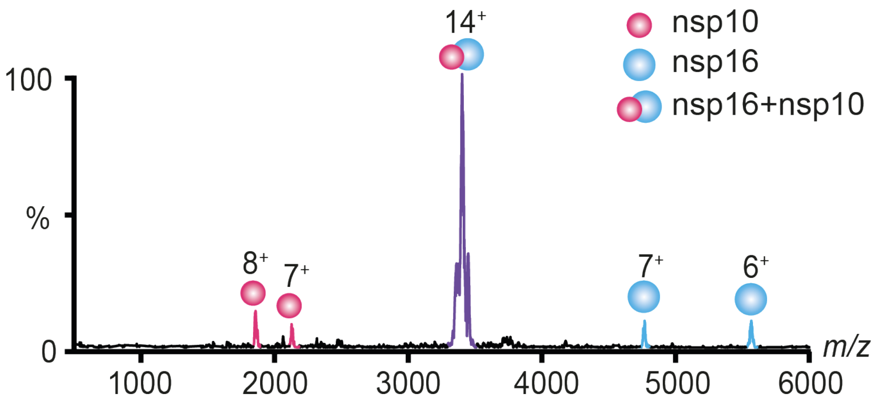


**Supplementary Figure S3: Tandem MS with CID on $14^+$-precursor ion of nsp16/10 (48.4 kDa) complex.**

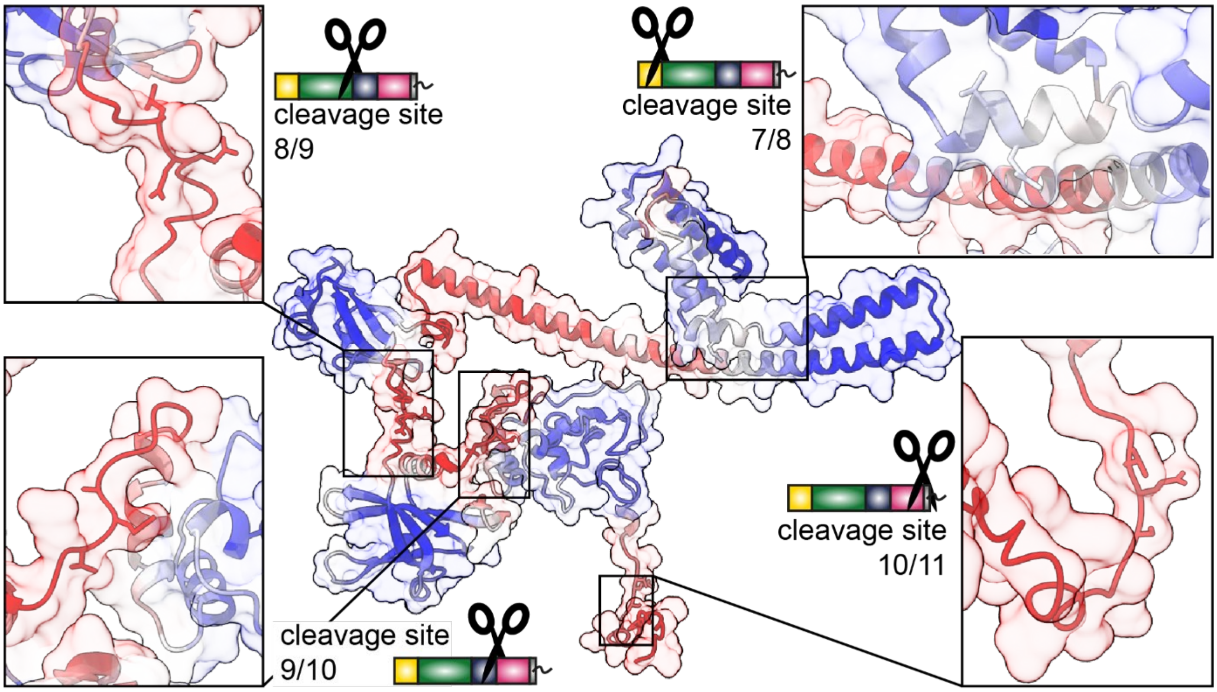


**Supplementary Figure S4: Alphafold2 model colored by *B*-factor using a palette from blue, white, to red with ranges set from 50 to 90.**

**Supplementary Table S1: Cleavage area is defined here as 5 residues before and after the corresponding cleavage sites LQ/A/S/N (P6-P6').** Confidence scores from Alphafold2 for the cleavage area were averaged.

| Cleavage area | Residues | Average confidence score |
|---|---|---|
| **7/8** | 77-89 | 73.5 |
| **8/9** | 275-287 | 34.9 |
| **9/10** | 388-400 | 39.6 |
| **10/11** | 527-539 | 40.9 |

**Supplementary Table S2: Amino acid sequences of all recombinantly expressed proteins and their theoretical mass (in Da)**

| Protein | Sequence | Theoretical Mass (in Da) |
|---|---|---|
| **nsp7-11C** | SKMSDVKCTSVVLLSVLQQLRVESSSKLWAQCVQLHND<br>ILLAKDTTEAFEKMVSLLSVLLSMQGAVDINKLCEEMLDN<br>RATLQAIASEFSSLPSYAAFATAQEAYEQAVANGDSEVV<br>LKKLKKSLNVAKSEFDRDAAMQRKLEKMADQAMTQMY<br>KQARSEDKRAKVTSAMQTMLFTMLRKLDNDALNNIINNA<br>RDGCVPLNIIPLTTAAKLMVVIPDYNTYKNTCDGTTFTYA<br>SALWEIQQVVDADSKIVQLSEISMDNSPNLAWPLIVTALR<br>ANSAVKLQNNELSPVALRQMSCAAGTTQTACTDDNALA<br>YYNTTKGGRFVLALLSDLQDLKWARFPKSDGTGTIYTEL<br>EPPCRFVTDTPKGPKVKYLYFIKGLNNLNRGMVLGSLAA<br>TVRLQAGNATEVPANSTVLSFCAFAVDAAKAYKDYLASG<br>GQPITNCVKMLCTHTGTGQAITVTPEANMDQESFGGAS<br>CCLYCRCHIDHPNPKGFCDLKGKYVQIPTTCANDPVGFT<br>LKNTVCTVCGMWKGYGCSCDQLREPMLQSADAQSFLN<br>GFAVSARGSHHHHHH | 60,824 |
| **nsp7-11N** | ASRGSHHHHHHGASKMSDVKCTSVVLLSVLQQLRVESS<br>SKLWAQCVQLHNDILLAKDTTEAFEKMVSLLSVLLSMQG<br>AVDINKLCEEMLDNRATLQAIASEFSSLPSYAAFATAQEA<br>YEQAVANGDSEVVLKKLKKSLNVAKSEFDRDAAMQRKL<br>EKMADQAMTQMYKQARSEDKRAKVTSAMQTMLFTMLR<br>KLDNDALNNIINNARDGCVPLNIIPLTTAAKLMVVIPDYNT<br>YKNTCDGTTFTYASALWEIQQVVDADSKIVQLSEISMDN<br>SPNLAWPLIVTALRANSAVKLQNNELSPVALRQMSCAAG<br>TTQTACTDDNALAYYNTTKGGRFVLALLSDLQDLKWAR<br>FPKSDGTGTIYTELEPPCRFVTDTPKGPKVKYLYFIKGLN<br>NLNRGMVLGSLAATVRLQAGNATEVPANSTVLSFCAFA<br>VDAAKAYKDYLASGGQPITNCVKMLCTHTGTGQAITVTP<br>EANMDQESFGGASCCLYCRCHIDHPNPKGFCDLKGKYV<br>QIPTTCANDPVGFTLKNTVCTVCGMWKGYGCSCDQLRE<br>PMLQSADAQSFLNGFAV | 60,953 |
| **nsp16** | MHHHHHHSAVLQSSQAWQPGVAMPNLYKMQRMLLEK<br>CDLQNYGDSATLPKGIMMNVAKYTQLCQYLNTLTLAVPY<br>NMRVIHFGAGSDKGVAPGTAVLRQWLPTGTLLVDSDLN<br>DFVSDADSTLIGDCATVHTANKWDLIISDMYDPKTKNVT<br>KENDSKEGFFTYICGFIQQKLALGGSVAIKITEHSWNADL<br>YKLMGHFAWWTAFVTNVNASSSEAFLIGCNYLGKPREQ<br>IDGYVMHANYIFWRNTNPIQLSSYSLFDMSKFPLKLRGT<br>AVMSLKEGQINDMILSLLSKGRLIIRENNRVVISSDVLVNN | 34,776 |
| **M$^{pro}$** | SGFRKMAFPSGKVEGCMVQVTCGTTTLNGLWLDDVVY<br>CPRHVICTSEDMLNPNYEDLLIRKSNHNFLVQAGNVQLR<br>VIGHSMQNCVLKLKVDTANPKTPKYKFVRIQPGQTFSVL<br>ACYNGSPSGVYQCAMRPNFTIKGSFLNGSCGSVGFNID<br>YDCVSFCYMHHMELPTGVHAGTDLEGNFYGPFVDRQT<br>AQAAGTDTTITVNVLAWLYAAVINGDRWFLNRFTTTLND<br>FNLVAMKYNYEPLTQDHVDILGPLSAQTGIAVLDMCASL<br>KELLQNGMNGRTILGSALLEDEFTPFDVVRQCSGVTF | 33,669 |

**Supplementary Table S3: All mass species that were detected on the Q Exactive UHMR Orbitrap. Measured masses and FWHM were averaged and standard error is given.**

| Name | Theoretical Mass (in Da) | Measured mass (in Da) | FWHM (in Da) |
|---|---|---|---|
| nsp7 | 9,239.8 | 9,239 ± 0.1 | 6.2 ± 0.1 |
| nsp7N | 10,649.3 | 10,649 ± 1 | 7 ± 1 |
| nsp9 | 12,378.2 | 12,378 ± 0.2 | 7.2 ± 0.1 |
| nsp10+$Zn_2$ | 14,919.9 | 14,916 ± 0.2 | 10 ± 1 |
| nsp10-11+$Zn_2$ | 17,509.7 | 17,506 ± 1 | 10 ± 1 |
| $nsp7_2$ | 18,479.6 | 18,479 ± 0.2 | 11 ± 1 |
| nsp8 | 21,881.1 | 21,881 ± 0.1 | 10.1 ± 0.1 |
| nsp9-10+$Zn_2$ | 27,280.1 | 27,276 ± 0.1 | 13.6 ± 0.2 |
| nsp9-11+$Zn_2$ | 29,869.8 | 29,866 ± 0.2 | 15 ± 1 |
| nsp7-8 | 31,102.9 | 31,103 ± 1 | 15 ± 1 |
| nsp7+8 | 31,120.9 | 31,120 ± 1 | 15 ± 1 |
| nsp7-8N | 32,512.3 | 32,512 ± 1 | 16 ± 2 |
| nsp7N+8 | 32,530.4 | 32,530 ± 1.2 | 17 ± 1 |
| $M^{pro}$ | 33,668.5 | 33,796 ± 1 | 16 ± 2 |
| nsp7-9 | 43,463.1 | 43,462 ± 1 | 23 ± 2 |
| $nsp8_2$ | 43,762.2 | 43,762 ± 2 | 26 ± 3 |
| nsp7-10C+$Zn_2$ | 58,365.0 | 58,360 ± 2 | 36 ± 2 |
| nsp7-10N+$Zn_2$ | 59,774.7 | 59,774 ± 3 | 41 ± 2 |
| nsp7-11C+$Zn_2$ | 60,954.7 | 60,950 ± 4 | 39 ± 2 |
| nsp7-11N+$Zn_2$ | 61,082.9 | 61,085 ± 1 | 42 ± 1 |
| $nsp7_2+8_2$ | 62,241.8 | 62,244 ± 5 | 38 ± 3 |
| nsp7N+7 +$8_2$ | 63,651.3 | 63,655 ± 3 | 41 ± 2 |
| $nsp7N_2+8_2$ | 65,024.6 | 65,061 ± 3 | 41 ± 2 |
| $M^{pro}_2$ | 67,337.0 | 67,598 ± 1 | 40 ± 1 |

**Supplementary Table S4: Masses for nsp16/10 interaction study as measured on a Q-TOF2 modified for high mass species.** Measured mass and FWHM was averaged.

| Name | Theoretical Mass (in Da) | Measured mass (in Da) | FWHM (in Da) |
|---|---|---|---|
| nsp9 | 12,378.2 | 12415 ± 11 | 82 ± 5 |
| nsp10+$Zn_2$ | 14,919.9 | 14951 ± 15 | 120 ± 52 |
| $nsp7_2$ | 18,479.64 | 18506 ± 6 | 110 ± 20 |
| nsp16 | 33,323.32 | 33390 ± 12 | 210 ± 20 |
| $M^{pro}$ | 33,668.51 | 33940 ± 40 | 250 ± 50 |
| nsp16-$His_6$ | 34,775.94 | 34890 ± 22 | 280 ± 90 |
| nsp16+10+$Zn_2$ | 48, | 48361 ± 5 | 280 ± 20 |
| nsp7-11C+$Zn_2$ | 60,954.7 | 61140 ± 18 | 410 ± 90 |
| $M^{pro}_2$ | 67,337.02 | 67840 ± 50 | 1000 ± 700 |

**Supplementary Table S5: Experimental determined masses (in Da) and list of gates that were used for the analysis of nsp711C.**

| Name | Mass | charge states | *m/z* min | *m/z* max |
|---|---|---|---|---|
| nsp10+$Zn_2$ | 14920 | 4 | 3728 | 3732 |
| nsp10+$Zn_2$ | 14920 | 5 | 2982 | 2985 |
| nsp10+$Zn_2$ | 14920 | 6 | 2485.5 | 2488 |
| nsp10+$Zn_2$ | 14920 | 7 | 2130.5 | 2133 |
| nsp10-11+$Zn_2$ | 17510 | 6 | 2916 | 2921 |
| nsp10-11+$Zn_2$ | 17510 | 7 | 2500 | 2503 |
| $M^{pro}{}_2$ | 67594 | 15 | 4504.5 | 4511 |
| $M^{pro}{}_2$ | 67594 | 16 | 4223 | 4230 |
| $M^{pro}{}_2$ | 67594 | 17 | 3974.5 | 3981 |
| $M^{pro}$ | 33797 | 10 | 3378 | 3383 |
| $M^{pro}$ | 33797 | 11 | 3071 | 3076 |
| $M^{pro}$ | 33797 | 12 | 2815.5 | 2819 |
| $M^{pro}$ | 33797 | 13 | 2599 | 2602 |
| nsp7-10+$Zn_2$ | 58365 | 13 | 4487.5 | 4493 |
| nsp7-10+$Zn_2$ | 58365 | 14 | 4167 | 4173 |
| nsp7-10+$Zn_2$ | 58365 | 17 | 3889.5 | 3895 |
| nsp7-11+$Zn_2$ | 60958 | 13 | 4687 | 4693 |
| nsp7-11+$Zn_2$ | 60958 | 14 | 4352 | 4358 |
| nsp7-11+$Zn_2$ | 60958 | 15 | 4062 | 4067 |
| nsp7-11+$Zn_2$ | 60958 | 16 | 3808 | 3812 |
| nsp7-8 | 31121 | 9 | 3454 | 3458.5 |
| nsp7-8 | 31121 | 10 | 3109.5 | 3113 |
| nsp7-8 | 31121 | 11 | 2826.5 | 2830 |
| $nsp7_2+8_2$ | 62242 | 14 | 4442 | 4450 |
| $nsp7_2+8_2$ | 62242 | 15 | 4146 | 4153 |
| nsp7-9 | 43463 | 12 | 3620.5 | 3626 |
| nsp7-9 | 43463 | 13 | 3342 | 3346 |
| nsp7-9 | 43463 | 14 | 3103.5 | 3107 |
| $nsp7_2$ | 18480 | 7 | 2639.5 | 2642 |
| $nsp7_2$ | 18480 | 9 | 2053 | 2055 |
| nsp7 | 9240 | 3 | 3078.5 | 3082.5 |
| nsp7 | 9240 | 4 | 2308.5 | 2312 |
| nsp7 | 9240 | 5 | 1847.5 | 1850 |
| $nsp8_2$ | 43762 | 10 | 4373.5 | 4379.5 |
| $nsp8_2$ | 43762 | 11 | 3974.5 | 3980.5 |
| $nsp8_2$ | 43762 | 12 | 3645 | 3649.5 |
| nsp8 | 21881 | 8 | 2734 | 2738 |
| nsp8 | 21881 | 9 | 2430.5 | 2433.5 |
| nsp9-10+$Zn_2$ | 27280 | 8 | 3408 | 3412 |
| nsp9-10+$Zn_2$ | 27280 | 9 | 3029.5 | 3034 |
| nsp9-11+$Zn_2$ | 29870 | 9 | 3317.5 | 3322 |
| nsp9-11+$Zn_2$ | 29870 | 10 | 2985.5 | 2990 |
| nsp9-11+$Zn_2$ | 29870 | 11 | 2714 | 2718 |
| nsp9 | 12378 | 4 | 3093 | 3097 |
| nsp9 | 12378 | 5 | 2474 | 2478 |

| | | | | |
|---|---|---|---|---|
| nsp9 | 12378 | 6 | 2062.5 | 2065 |
| nsp9 | 12378 | 7 | 1768 | 1770 |

**Supplementary Table S6: List of masses measured and gates that were used for the analysis nsp7 11N.**

| Name | Mass | charge states | *m/z* min | *m/z* max |
|---|---|---|---|---|
| nsp7N | 10650 | 3 | 3078.5 | 3084 |
| nsp7N | 10650 | 4 | 2661 | 2665 |
| nsp7N | 10650 | 5 | 2129 | 2132 |
| nsp7-8N | 32512 | 9 | 3611 | 3616 |
| nsp7-8N | 32512 | 10 | 3250 | 3256 |
| nsp7-8N | 32512 | 11 | 2955 | 2958 |
| nsp7-8 | 31121 | 9 | 3454 | 3458.5 |
| nsp7-8 | 31121 | 10 | 3109.5 | 3113 |
| nsp7-8 | 31121 | 11 | 2826.5 | 2830 |
| nsp7-10N | 59780 | 13 | 4595 | 4604 |
| nsp7-10N | 59780 | 14 | 4267 | 4274 |
| nsp7-11N | 61090 | 13 | 4696 | 4704 |
| nsp7-11N | 61090 | 14 | 4361 | 4368 |
| nsp7-11N | 61090 | 15 | 4070 | 4075 |
| nsp9-10 | 27280 | 8 | 3408 | 3412 |
| nsp9-10 | 27298 | 9 | 3029.5 | 3034 |
| nsp9-10 | 27298 | 10 | 2726.5 | 2730.5 |
| nsp7 | 9240 | 5 | 1847.5 | 1850 |
| nsp7 | 9240 | 4 | 2308.5 | 2312 |
| nsp7 | 9240 | 3 | 3078.5 | 3082.5 |
| $nsp7_2$ | 18480 | 9 | 2053 | 2055 |
| $nsp7_2$ | 18480 | 7 | 2639.5 | 2642 |
| nsp8 | 21881 | 9 | 2430.5 | 2433.5 |
| nsp8 | 21881 | 8 | 2734 | 2738 |
| $nsp8_2$ | 43762 | 12 | 3645 | 3649.5 |
| $nsp8_2$ | 43762 | 11 | 3974.5 | 3980.5 |
| $nsp8_2$ | 43762 | 10 | 4373.5 | 4379.5 |
| $nsp7_2+8_2$ | 62242 | 15 | 4146 | 4153 |
| $nsp7_2+8_2$ | 62242 | 14 | 4442 | 4450 |
| nsp9 | 12378 | 7 | 1768 | 1770 |
| nsp9 | 12378 | 6 | 2062.5 | 2065 |
| nsp9 | 12378 | 5 | 2474 | 2478 |
| nsp9 | 12378 | 4 | 3093 | 3097 |
| nsp10 | 14920 | 4 | 3728 | 3732 |
| nsp10 | 14920 | 7 | 2130.5 | 2133 |
| nsp10 | 14920 | 6 | 2485.5 | 2488 |
| nsp10 | 14920 | 5 | 2982 | 2985 |
